# Q-FADD: A mechanistic approach for modeling the accumulation of proteins at sites of DNA damage by free diffusion

**DOI:** 10.1101/373043

**Authors:** Jyothi Mahadevan, Johannes Rudolph, Asmita Jha, Jian Wei Tay, Joe Dragavon, Erik M. Grumstrup, Karolin Luger

## Abstract

The repair of DNA damage requires the ordered recruitment of many different proteins that are responsible for signaling and subsequent repair. A powerful tool for studying the orchestrated accumulation of these proteins at damage sites is laser microirradiation in live cells, followed by monitoring of the accumulation of the fluorescently labeled protein in question. Despite the widespread use of this approach, there exists no rigorous method for characterizing this process quantitatively. Here we introduce a free diffusion model that explicitly accounts for the unique topology of individual nuclei and quantitatively describes the accumulation of two test proteins, poly-ADP-ribose polymerases 1 and 2. Application of our model to other proteins will yield novel insights into the timing and mechanism of DNA repair.

## Introduction

Genomic DNA is continuously subjected to endogenous and exogenous insults from free radicals, ionizing radiation, and DNA-modifying chemicals. All of these, either directly or during the repair process, cause single-strand and double-strand breaks (SSBs and DSBs). Without proper DNA damage detection and repair, the resulting genomic instabilities can lead to premature aging, sensitivity toward radiation damage, and cancer. While specific repair pathways exist for different types of DNA lesions, the sequential accumulation and binding of many signaling and repair proteins at damage sites is conserved^1^. Thus, a quantitative understanding of protein accumulation, order of accumulation, and variability due to type and amount of damage, cell type, etc. is critical not only for establishing a fundamental framework of DNA repair pathways, but is also of clinical relevance as mutations or mis-regulation in DNA damage detection and repair pathways are strongly associated with, or even cause many cancers^2^.

There are many different genetic, biochemical, cellular, and animal-model methods to investigate the various DNA repair pathways. Laser microirradiation is a particularly powerful method to study these processes in living cells. In this approach, cells are first transiently transfected with a protein of interest that has been fluorescently tagged. Local DNA damage is induced in the transfected cells with a short wavelength (355-405 nm) confocal laser beam, after which the time-dependent accumulation of the proteins of interest at the site of laser-induced DNA damage is monitored with fluorescence microscopy. This unique combination of biophysical manipulation and cell biology has yielded compelling data on the order and kinetics for the recruitment of many different DNA repair proteins such as PARP1, ATM kinase, Ku80, p21, and even phospholipids at sites of DNA damage ^3,4,13,5–12^.

While extensive analytical methods have been developed for quantitation of Fluorescence Recovery After Photobleaching (FRAP) and Fluorescence Loss In Photobleaching (FLIP)^14–16^, analysis of laser microirradiation experiments typically is limited to reports of simple appearance at sites of damage, usually expressed as the time to half-maximal accumulation (t_1/2_)^10^. Fitting to multiple first order rate constants that have no physical basis is also done occasionally^3^. Such methods may be useful for determining the relative order of accumulation within a single cell, but they do not provide any insight into the mechanism of transport, such as simple, facilitated, or anomalous diffusion. Furthermore, these analyses ignore the effects of variable size and shape of different nuclei, which, as we show here, can lead to qualitatively and quantitatively incorrect conclusions. Here, we start with the simplest assumption that PARPs, like most nuclear proteins, move by simple diffusion. To test this, we introduce a Monte Carlo free diffusion model that explicitly accounts for the unique topology of each nucleus. The model, Q-FADD (<u>Q</u>uantitation of <u>F</u>luorescence <u>A</u>ccumulation after <u>D</u>NA <u>D</u>amage), suitably describes the accumulation kinetics of PARP1 and PARP2 at sites of DNA damage, leading to novel mechanistic insights into the properties of these DNA repair proteins.

## Results

### Laser Microirradiation and Accumulation of PARPs at Sites of DNA Damage

We transfected either GFP-PARP1 or GFP-PARP2 into mouse embryo fibroblasts (MEFs) where both proteins were easily visualized in nuclei by fluorescence microscopy (e.g. Figure 1A). We next monitored the recruitment of GFP-PARP1 and GFP-PARP2 to sites of laser irradiation (see Figure 1A; representative movies can be found in SuppMovie1 and SuppMovie2). In agreement with previous reports^8,17^, we saw an increase in fluorescence intensity of PARP1 and PARP2 at the site of DNA damage by factors of 1.2 to 5.5, reaching maximum intensity within 60 – 200 s (Figure 1B). As noted previously^8^, PARP1 accumulates more rapidly than PARP2, as measured by t_1/2_ of averaged nuclei (Fig 1B). For both PARP1 and PARP2, we observed significant depletion of fluorescence from sites outside the region of DNA damage (Figure 1A), and as reported previously, both PARP1 and PARP2 deplete from the site of damage when monitored for much longer time courses (not shown)^8,17^. To quantify the difference in rates of accumulation between PARP1 and PARP2, we began by attempting to fit our data from individual nuclei using single or multiple exponentials, as is the standard in the literature. This approach yielded highly variable results; and moreover, it has no physical basis. We thus turned to developing a method based on the physics of free diffusion.

**Figure 1:**
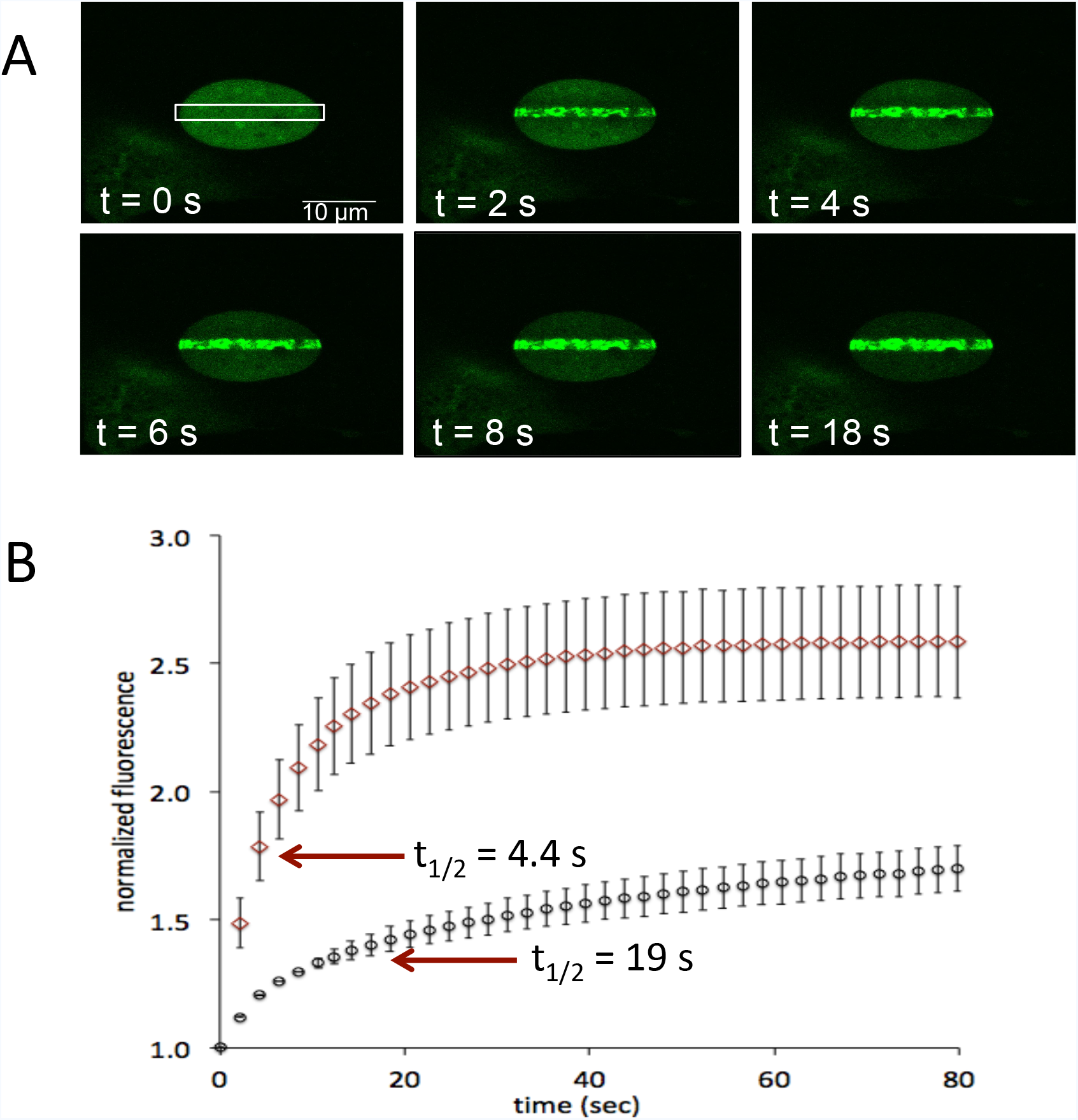
PARP1 accumulates faster than PARP2 at sites of DNA damage after laser microirradiation. A) Movie snapshots showing GFP-PARP1 accumulating at a site of DNA damage (white box) in a MEF nucleus. B) Accumulation of GFP-PARP1 (diamonds) and GFP-PARP2 (circles) in MEF cells shown as the average ± SEM of 28 and 19 nuclei, respectively. Accumulation of PARP1 and PARP2 for individual nuclei are shown in Supp. Figure 4 and Supp. Figure 5, respectively.

### Modeling of Free Diffusion

While a continuum diffusion approach could in principle be used to model experimental results (as is the standard for FRAP^14^), the boundary conditions for each uniquely shaped nucleus would be challenging to parameterize. We therefore modeled protein diffusion using a Monte Carlo approach, which provides the ability to simulate a discrete diffusional process while incorporating an experimentally accurate nuclear profile into the simulation. In these simulations, the diffusion coefficient (D_eff_) of the particles is parameterized with the particle step frequency (1/Δt) and the particle step size (Δx). The trajectories of the particles as they undergo a random two-dimensional walk are confined to the experimentally accurate nuclear boundary and are trapped at the defined site of laser damage.

Nuclei in living cells are highly variable in topology (see e.g. Supp. Figs. 4 - 7). To highlight the importance of taking into account both the size and shape of a nucleus to correctly determine D_eff_, we modeled a series of simple test cases, which are presented in Figure 2. In the simplest example, we model diffusion using the same D_eff_ in three different circular nuclei of varying diameter (Figure 2A). Apparent accumulation as measured by t_1/2_ is slowest in the largest circle, consistent with the intuition that the average molecule must traverse a longer distance to be caught in the trap. Next, we keep both D_eff_ and the area of each nucleus constant but vary the ellipticity (Figure 2B). As measured by t_1/2_, the accumulation of particles takes longer in more elongated nuclei, again consistent with the intuition that the average molecule must traverse a longer distance to be caught in the trap. In a final example, which most realistically mimics the diversity we observe in real nuclei, we allow both the size and ellipticity to vary, while holding constant the size of the trap and D_eff_ (Figure 2C). Again, particles appear to accumulate more slowly in the more elliptical and larger nuclei. We emphasize that in all three examples, D_eff_ is constant, yet the t_1/2_ parameters vary by as much as a factor of 10 because the size and shape of the nuclei are changing.

**Figure 2:**
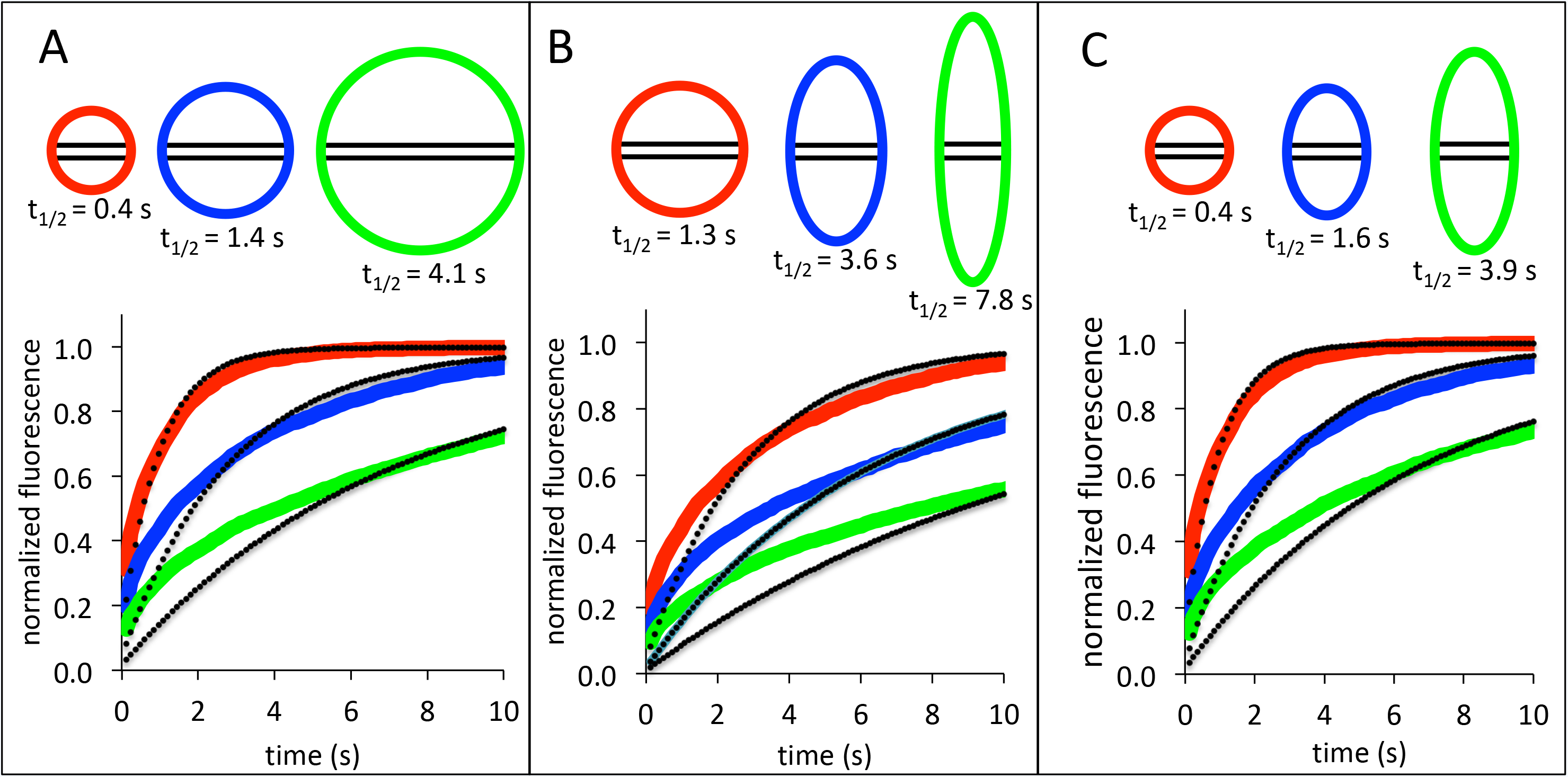
Varying nucleus size and shape affect t_1/2_ at constant values of D_eff_. Each panel shows a different scenario of size and/or shape variation of the nucleus along with the simulated accumulation of GFP-PARP1 in the region of laser damage (box, with unchanged height), and the apparent t_1/2_ assuming a D_eff_ = 4.53 μ^2^/s. The colors of the nuclei correspond to the data in the graph. A) Three different circular nuclei with radii of 60, 100, and 150 pixels. B) Three different nuclei of varying ellipticity (100×100, 70×143, 50×200 pixels) but with the same overall area. C) Three different nuclei of varying size and ellipticity (60×60, 60×100, 60×150 pixels), but, as in A) and B) with the same trap size. The black dots superimposed on each curve indicate the best-fit to a first order exponential, and demonstrate that size and shape also influence the quality of this fit, which may lead to arbitrary application of multi-exponential fitting for larger or more elongated nuclei.

Analysis of the accumulation kinetics using simple rate models leads to similarly misleading conclusions. For example, fits to the accumulation kinetics in Fig. 2 using a single exponential model (black dots) yield slower apparent derived rates for larger or more elongated nuclei compared to smaller and rounder nuclei, despite identical diffusion coefficients. Furthermore, as can be seen by comparing the single exponential fits in Figure 2C, larger or more elongated nuclei are fit more poorly to a single exponential than smaller rounder ones. Efforts to obtain better fits have in the past led to over-parameterization^3^. Thus, unless applied only as a comparative tool in the *same nucleus*, analysis of protein accumulation at sites of DNA damage by either t_1/2_ measurements or exponential models will lead to specious results, which are both qualitatively and quantitatively incorrect.

### Testing the Free Diffusion Model for PARP1 and PARP2 recruitment

In order to quantify the difference in rates of accumulation between PARP1 and PARP2, we used our model of free diffusion to simulate the accumulation of PARPs in the region of DNA damage in MEF cells, comparing the modeled curves with actual experimental data. For both PARP1 and PARP2, two parameters were sufficient to generate curves that fit the data (as judged by r-squared coefficient >0.96), namely D_eff_, the coefficient of free diffusion, and F, the fraction of mobile PARP. Three example datasets for PARP1 are shown in Figure 3A. In addition to the accumulation kinetics, our model also accurately describes the depletion of PARP1 from regions adjacent to the damage site using the same parameter values D_eff_ and F. We find that depletion occurs more quickly at sites closer to the damage site (Figure 3B). In Supp. Figure 4-5, we show the raw data, respective fits, and variability in size and shape for 28 nuclei accumulating GFP-PARP1, and 19 nuclei accumulating GFP-PARP2 at sites of DNA damage in MEF cells. We summarize the derived values of D_eff_ and F in box-and-whisker plots (Figure 4), and in Table 1.

**Figure 3:**
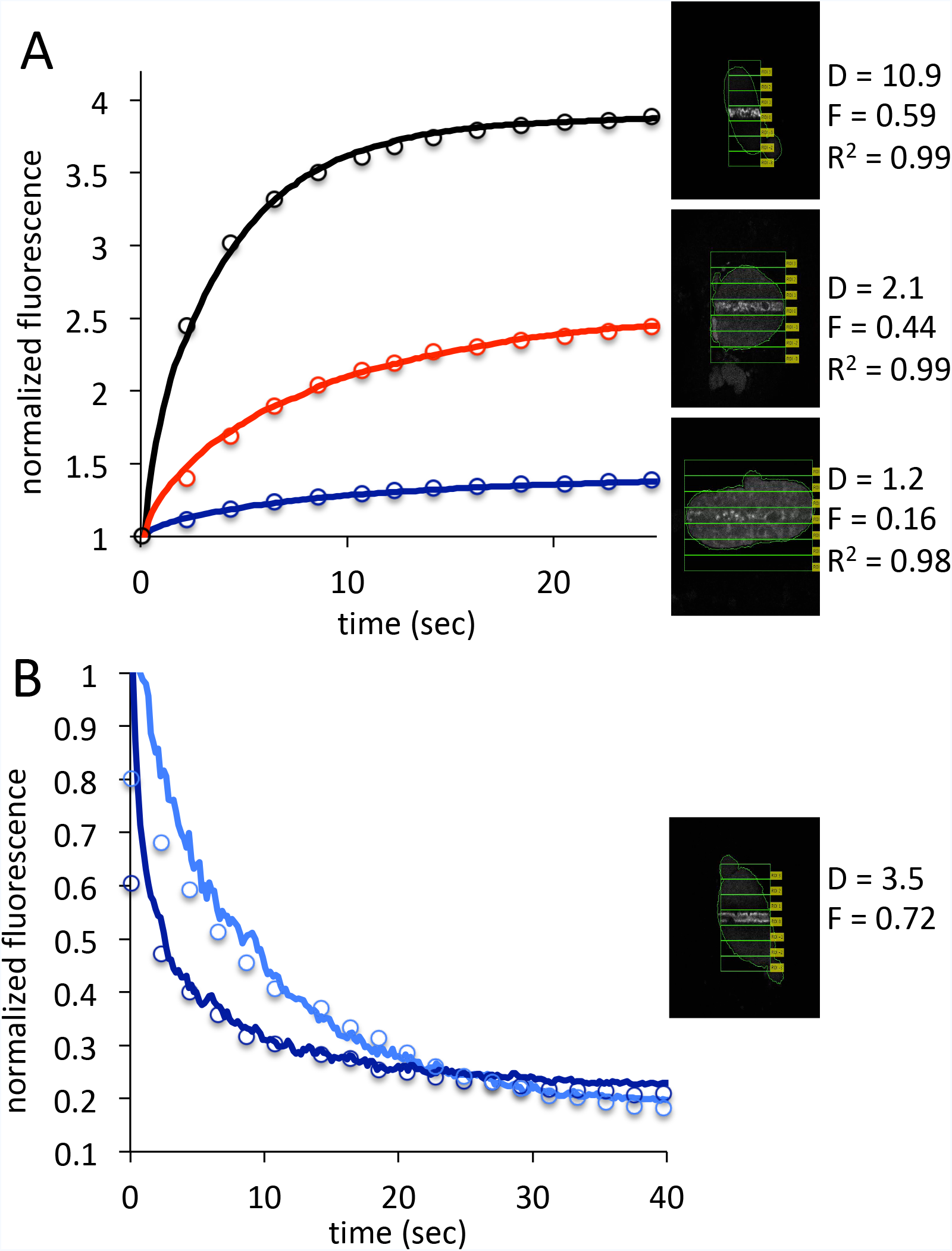
Both appearance of fluorescence at sites of DNA damage, and disappearance of fluorescence from the nucleus can be modeled by simple diffusion. A) Overlay of experimental data from Q-FADD for GFP-PARP1 and simulation to determine D_eff_ for GFP-PARP1 in MEF cells, for three representative nuclei (see also Supp. Figure 4). B) Depletion of PARP1 from two different sites within one nucleus can be quantitatively described using the same D_eff_ (dark blue = ROI-1, adjacent to site of damage, light blue = ROI-2, adjacent to ROI-1 (see also Supp Figure 2).

**Figure 4:**
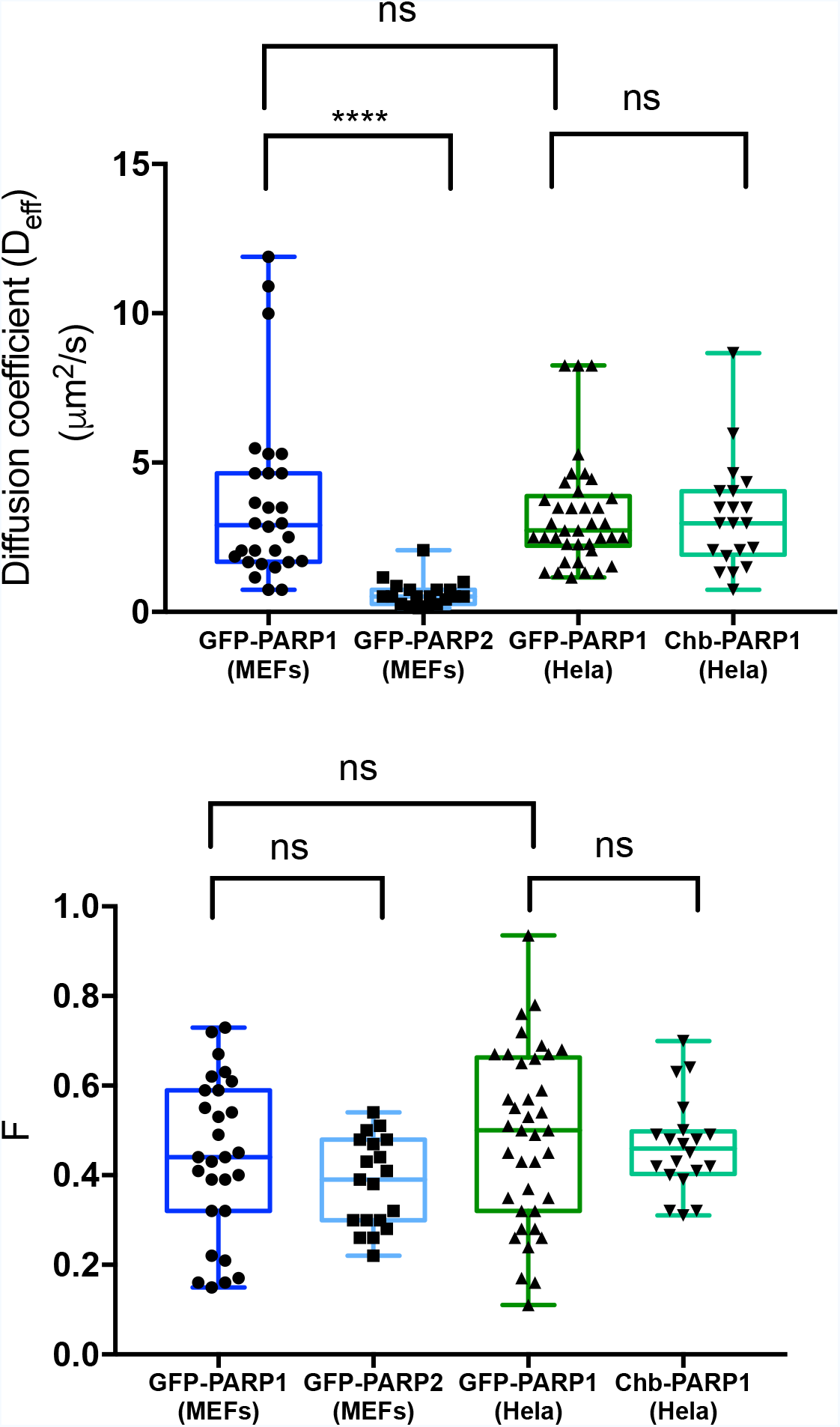
Q-FADD reveals statistically significant differences between PARP1 and PARP2 recruitment. Box and whisker plots for D_eff_ (in A) and F (in B) as determined by simulation and matching to Q-FADD with GFP-PARP1 and GFP-PARP2. Data are shown for transient transfections of GFP-PARP1 and GFP-PARP2 in MEF and HeLa cells, and detection of endogenous PARP1 using chromobodies (Chb). Statistical differences between pairs of samples were evaluated using the unpaired t-test with p < 0.001 (****).

**Table 1:** Summary of parameters derived from free diffusion analysis of Q-FADD data

|  | GFP-PARP1 | GFP-PARP2 | GFP-PARP1 | Chb-PARP1 |
| --- | --- | --- | --- | --- |
| Cell type | MEFs | MEFs | HeLa | HeLa |
| n | 28 | 19 | 38 | 20 |
| $D_{\text{eff}}$ ($\mu\text{m}^2/\text{s}$) ± SEM | $3.7 \pm 0.55$ | $0.62 \pm 0.1$ | $3.2 \pm 0.30$ | $3.2 \pm 0.41$ |
| F ± SEM | $0.44 \pm 0.03$ | $0.38 \pm 0.02$ | $0.48 \pm 0.03$ | $0.46 \pm 0.02$ |

Statistical analysis demonstrates that PARP2 has a six-fold slower diffusion coefficient than PARP1 at sites of DNA damage (Figure 4, Table 1). Test simulations show that this cannot be reproduced by assuming, based on experimental observations^18^, that PARP2 has a weaker affinity to DNA in the damaged site than PARP1. In fact, the opposite behavior is observed in simulations, with accumulation to maximum occurring faster when trapped particles are allowed to become free with finite probability.

We next investigated the effects of cell-type on the accumulation of PARP1 at sites of DNA damage. We monitored accumulation of GFP-PARP1 in HeLa cells with analysis by our free diffusion model and found statistically identical values for both D_eff_ and F compared to MEFs (Supp. Figure 6, Figure 4, and Table 1). To test the effects of endogenous vs. exogenous PARP1 expression, we needed to find a way to measure the accumulation of endogenous (untagged) PARP1. Chromobodies are small functional antibodies that are tagged with a chromophore and are readily transfected into cells^19^. Chromobodies to human PARP1 were used to monitor PARP1 accumulation in HeLa cells, to compare with accumulation of transiently transfected GFP-PARP1. Subsequent analysis also yielded statistically identical values for both D_eff_ and F as confirmed by unpaired Student’s t test (Supp. Figure 7, Figure 4, and Table 1). These results demonstrate that transfection of GFP-tagged PARP1 does not significantly affect its accumulation at sites of DNA damage in the nucleus.

### Correlation analysis

Our extensive description of the variability between different nuclei in D_eff_ and F for PARP1 and PARP2 is novel in the analysis of protein accumulation by laser microirradiation, where like in FRAP and FLIP, data are generally presented as averages of many different nuclei. Interestingly, both D_eff_ and F varied significantly for different nuclei, despite taking into account the different sizes and shapes of the nuclei (note the whiskers in Figure 4). Using our largest data set of 38 nuclei of GFP-PARP1 in HeLa cells, we were unable to correlate D_eff_ or F with the initial fluorescence (Supp. Figure 8), suggesting that natural variability in expression levels of fluorescent PARP1 does not cause a change in mobility. Thus, Q-FADD is not biased towards nuclei with high or low initial fluorescence, alleviating the need of imaging only those nuclei that have similar initial fluorescence values. Additionally, we saw no correlation of D_eff_ or F with initial levels of signal from Hoechst 33342 (Supp Figure 8), which is a measure of a complex combination of DNA content, chromatin condensation, and/or apoptotic state.

Most interestingly, we found that there was no correlation between D_eff_ and the fraction of mobile protein, F (Figure 5). In analysis of diffusion by FRAP, F is typically interpreted to be the ratio of the effective rates of protein binding and release from stationary sites (i.e. DNA)^14^. If this assumption were to hold for PARP1, we would expect a direct correlation between D_eff_ and F, as a larger F would imply less binding of PARP1 during its travel from its initial position to the damage site, and thus faster arrival and a larger observed D_eff_. The fact that we do not observe a correlation between D_eff_ and F strongly suggests that this assumption is incorrect, at least for PARP1 and PARP2 (data for PARP2 not shown). The lack of correlation and the variability in both D_eff_ and F for PARP1 and PARP2 hint at a yet unidentified cause for the cell-to-cell variability that is worthy of further investigation. For example, one might expect to see variable rates of free diffusion depending on cell cycle state or levels of DNA damage, and this is an area of future studies.

**Figure 5:**
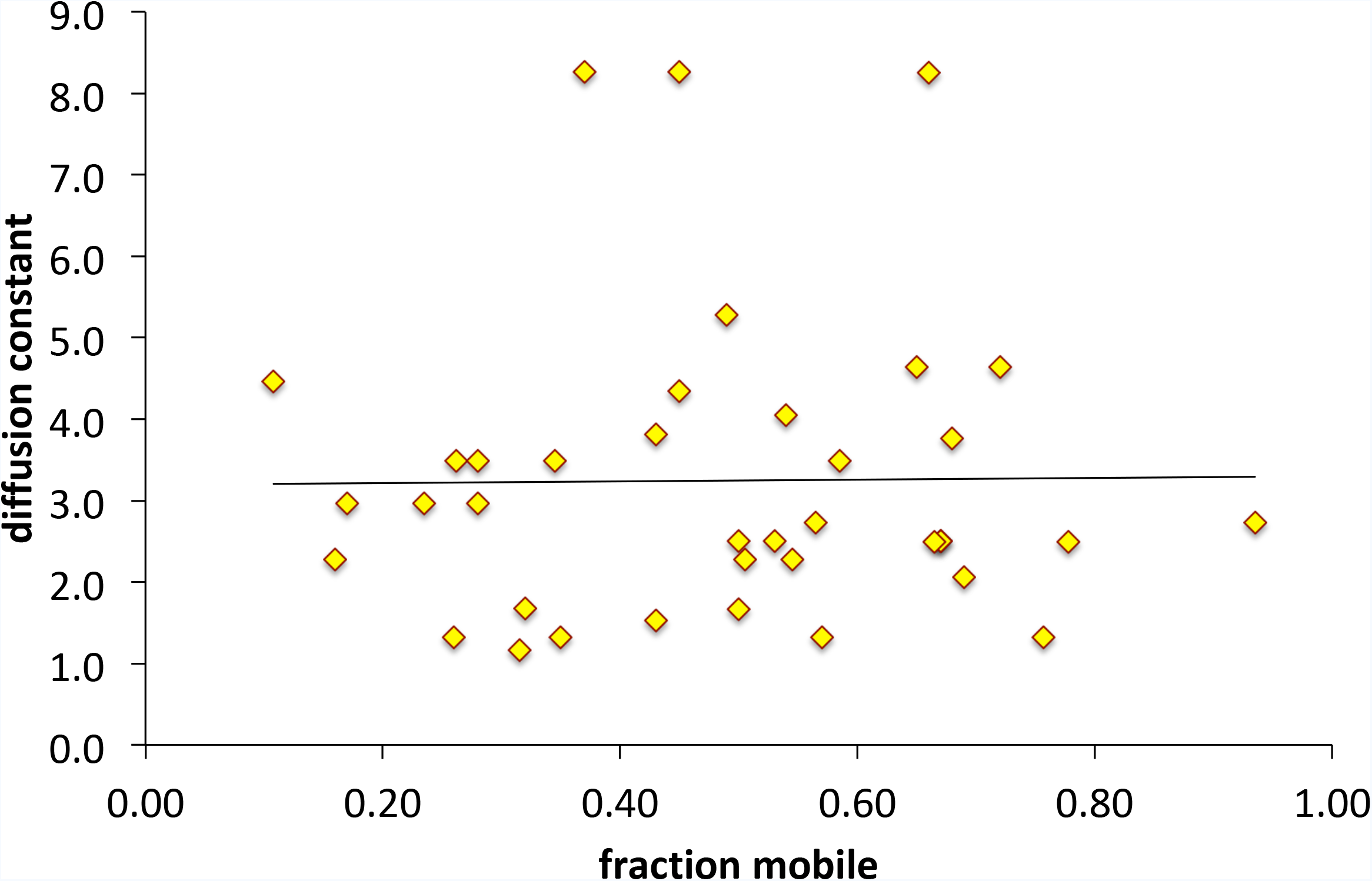
D_eff_ and F are not correlated. D_eff_ is plotted vs F for the accumulation kinetics of 38 HeLa nuclei transfected with GFP-PARP1 and analyzed by Q-FADD. A similar lack of correlation was seen for GFP-PARP2 (data not shown).

## Discussion

For PARP1 and PARP2, accumulation at sites of DNA damage can be described by simple free diffusion. Unlike some other proteins, our results imply that PARP1 and PARP2 movement is neither facilitated (i.e. actively transported by directed motion) nor anomalous^14–16, 20^. The mean D_eff_ of GFP-PARP1 is very fast (Table 1), only ~9-fold slower than the theoretical limit of 31.5 μm^2^/s; assuming a spherical protein of 143 kDa (r = 3.45 nm) and a viscosity of the nuclear milieu of 2 ×10^−3^ N s/m^2^, two-fold as viscous as pure water^21^. In fact, the D_eff_ of PARP1 is among the fastest known of any nuclear protein whose D_eff_ have been determined by FRAP, FLIP, or single molecule tracking^14,22,23^. The high D_eff_ for PARP1 is surprising since PARP1 is known to bind tightly to damaged <u>and</u> undamaged DNA^24,25^, and in light of the fact that PARP1 has many other roles that involve binding to chromatin or other proteins^26^. The fast diffusion of PARP1 is consistent with our recent demonstration of DNA release from PARP1 being facilitated by binding of an additional strand of DNA (monkey-bar mechanism), which is in high abundance throughout the nucleus^27^. Although PARP2 is smaller than PARP1, and thereby expected to move more rapidly, GFP-PARP2 diffuses slower than GFP-PARP1. In fact, it diffuses 60-fold slower than its theoretical limit of 36.4 μm^2^/s. The accumulation kinetics of PARP2 do not show a characteristic lag, a feature that has been previously attributed to sequential accumulation^3^, i.e. PARP2 recruitment does not depend on accumulation of some other protein, such as PARP1. Perhaps PARP2, which unlike PARP1 only has one DNA-binding domain^28^, does not use the monkey-bar mechanism to facilitate its movement inside the nucleus. Although the reason for the slower diffusion of PARP2 is unknown at this time, the application of Q-FADD now allows for quantitative investigations using mutants of PARP2 and/or knock-downs of potential binding partners.

In summary, we have presented a powerful new method that combines the cell-based technique of laser-induced microirradiation with Monte Carlo simulation to derive the diffusion coefficient D_eff_ and the fraction accumulation for quantifying proteins at sites of DNA damage. Because monitoring accumulation of fluorescent proteins following laser microirradiation is a relatively simple technique available to many researchers with standard cell culture and microscopy facilities, we anticipate that Q-FADD will find wide application in the field of DNA repair biology.

## Acknowledgements

We thank Francoise Dantzer (University of Strasbourg) for GFP-PARP clones. JM, JR, AJ, and KL are supported by NIH-NCI R01 CA218255, by the University of Colorado Cancer Center Pilot Funding Grant ST63501792, and the Howard Hughes Medical Institute. The imaging work was performed at the BioFrontiers Institute Advanced Light Microscopy Core. Laser scanning confocal microscopy was performed on a Nikon A1R microscope supported by NIST-CU Cooperative Agreement award number 70NANB15H226.

## Supplementary Figure Legends

Supp. Figure 1: (a) Greyscale image of the EGFP channel. Yellow circle indicates region used to calculate the intensity threshold. (b) The initial nuclear mask after calculating the intensity threshold. (c) The final nuclear mask after removing small objects, filling in gaps, and morphological dilation.

Supp. Figure 2: The white boxes correspond to the ROIs, while the outline of the nucleus is shown in green.

Supp. Figure 3: (a) The intensity histogram of the image was used to determine the background threshold level. The inset shows the full histogram, and the red box indicates the enlarged region. (b) The background (cyan) of each image was segmented and the mean intensity value was used for background subtraction for each ROI.

Supp. Figure 4: Overlay plots showing experimental data from Q-FADD with the simulation of free diffusion used to determine D_eff_ and F for GFP-PARP1 in MEF cells. For each of the 28 different nuclei, the values of D (in μm^2^/s), F, and R^2^ (r-squared coefficient) for each plot are shown along with a snapshot from each nucleus taken in the first frame after laser irradiation.

Supp. Figure 5: Overlay plots showing experimental data from Q-FADD with the simulation of free diffusion used to determine D_eff_ and F for GFP-PARP2 in MEF cells. For each of the 19 different nuclei, the values of D (in μm^2^/s), F, and R^2^ (r-squared coefficient) for each plot are shown along with a snapshot from each nucleus taken in the first frame after laser irradiation.

Supp. Figure 6: Overlay plots showing experimental data from Q-FADD with the simulation of free diffusion used to determine D_eff_ and F for GFP-PARP1 in HeLa cells. For each of the 38 different nuclei, the values of D (in μm^2^/s), F, and R^2^ (r-squared coefficient) for each plot are shown along with a snapshot from each nucleus taken in the first frame after laser irradiation.

Supp. Figure 7: Overlay plots showing experimental data from Q-FADD with the simulation of free diffusion used to determine D_eff_ and F for chromobody experiments detecting accumulation of endogenous PARP1 in HeLa cells. For each of the 20 different nuclei, the values of D (in μm^2^/s), F, and R^2^ (r-squared coefficient) for each plot are shown along with a snapshot from each nucleus taken in the first frame after laser irradiation.

Supp. Figure 8: There is no correlation between initial levels of GFP-PARP1 or initial staining by Hoechst 33342 with either D_eff_ or F for 38 HeLa nuclei. A) Initial fluorescent signal of GFP-PARP1 vs. D_eff_; B) Initial fluorescent signal of Hoechst 33342 vs. F. C) A) Initial fluorescent signal of GFP-PARP1 vs. F; D) Initial fluorescent signal of Hoechst 33342 vs. D_eff_. All fluorescent signals were divided by 10^6^. A similar lack of correlation was seen for GFP-PARP2 (data not shown).

Supp. Movie 1: Representative movie showing the accumulation of GFP-PARP1 at the site of DNA damage. This nucleus has a D_eff_ of 3.7 μ^2^/s and an F of 0.39.

Supp. Movie 2: Representative movie showing the accumulation of GFP-PARP2 at the site of DNA damage. This nucleus has a D_eff_ of 0.5 μ^2^/s and an F of 0.41.

## Materials and Methods

### Expression plasmids, cell culture and transient transfection

Mammalian expression plasmid (pEGFP-C3) encoding full-length GFP-tagged human PARP1 and wild type mouse embryo fibroblasts (MEFs) were a kind gift from Dr. Françoise Dantzer (University of Strasbourg). The expression plasmid for GFP-PARP2 was generated by subcloning human PARP2 cDNA into pEGFP-C3 between the restriction sites SalI and BamHI. PARP1 chromobody (Chb-PARP1) was purchased from ChromoTek GmbH (Germany).

MEF cells were cultured in DMEM supplemented with 50 μg/ml of gentamicin and 10% FBS. Hela-CCL2 cells were purchased from ATCC and were cultured in DMEM supplemented with 1X Penicillin/Streptomycin and 10% FBS. For the laser microirradiation experiments, cells were grown on CELLview TM slides (Greiner Bio-one) and were transfected with jetPEI (Polyplus Transfection) according to the manufacturer’s instructions. Briefly, 20,000 cells were plated and transfected 24 h later with 250 ng of DNA. Cells were sensitized with Hoechst 33342 (Invitrogen) (10μg/ml) for 10 min prior to the start of the experiment.

### Data collection

Live cell imaging and laser microirradiation experiments were performed on a Nikon A1R laser scanning confocal microscope equipped with a 405 nm diode laser, and an Argon-ion laser for 488 nm excitation (Biofrontiers Institute, University of Colorado, Boulder). Cells were imaged using a 100x 1.45NA oil immersion objective. A stage-top incubator was used to maintain proper environmental conditions of 37°C and 5% CO_2_, and for each data collection a cell was positioned so that its nucleus was at the center of the acquisition field of view. To induce DNA damage, the 405 nm laser was used at 100% power to irradiate a pre-determined rectangular region of interest (ROI) across the nucleus for 1s. Excitation of GFP fluorophore, using the 488 nm laser lines, was used to monitor the accumulation of PARP in the ROI. Six pre-irradiation and 150 post-irradiation frames were recorded at 2s time intervals. Images were collected at a frame size of 512 × 512 as per Nyquist sampling (1.2 AiryUnits). The resulting time lapse movie was saved as an ND2 file (proprietary Nikon format), while the region of irradiation was saved as a TIF image.

### Data Processing

Automated analysis of the fluorescent image was carried out using a custom code in MATLAB, which is described here. First, the ND2 image was imported into MATLAB using the Bioformats Image toolbox^29^. The region of the image corresponding to the nucleus was segmented using an intensity threshold from either the EGFP channel as converted to greyscale. The mean intensity *I_mean_* in a 20-pixel radius circle around the center of the image, i.e. the center of the nucleus (Supp. Figure 1a), was used to determine the threshold level (*I_mean_* – 0.5 × *I_mean_*). Applying this greyscale threshold to the image generated a binary mask of the nucleus (Supp. Figure 1b). Objects in the mask less than 500 pixels in area were removed and gaps in the mask were filled in. Finally, a disk-shaped structuring element (radius = 3 pixels) was used to perform morphological dilation and generate the final nuclear mask (Supp. Figure 1c). If multiple nuclei were present in the field-of-view, the nucleus closest to the center of the image was selected for further analysis.

To measure the fluorescence intensity across the nucleus as identified above, we divided the cell vertically into a number of region-of-interests (ROIs). The initial ROI was generated from the TIF image of the irradiation region and was vertically expanded by ten pixels to avoid edge effects. Depending on the height of the nucleus, additional ROIs were then generated by translating the original ROI above and below its original position (Supp. Figure 2). The total intensity of the nucleus within each ROI was measured for each frame of the movie. These intensity measurements were corrected for both background fluorescence and photobleaching. To determine the background fluorescence level, the image background was segmented using an intensity threshold. An intensity histogram of the image, smoothed by a 3-pixel moving window to remove spurious peaks, was used to set the threshold level. The first peak of the histogram, corresponding to the darkest greyscale values was identified and the greyscale value where the counts dropped to 1/e of the peak height was used to generate a mask of the background. The background intensity was then calculated as the mean intensity of the background mask, as shown in Supp Figure 3. This process was repeated for the first six pre-irradiation frames and the background correction level was calculated as the mean of these six values. This correction level was then subtracted for each frame of the movie. A separate correction level was calculated for each fluorescence channel in the movie. To correct for photobleaching, the total intensity of the nucleus in each frame was normalized to the total intensity of the nucleus in the first frame after applying the background correction. The correction for photobleaching was minimal (<10%).

For each ND2 image file, the code generated a CSV file containing the time series measurements of the intensity within each ROI. The ROI and the nuclear mask were also exported as text file to serve as inputs for the simulation of free diffusion.

### Simulation of Free Diffusion

Protein diffusion was modeled via standard Monte Carlo approaches on a two-dimensional grid using a custom code in the Mathematica environment. The region confined by the nuclear mask served as the active region of the simulation. 12,000 sample points were randomly generated at uniform density within the nuclear mask and allowed to propagate in a random walk. Control studies were performed with more sample points (up to 16,000), however results did not differ beyond a negligible reduction in noise observed for the accumulation and depletion kinetics. At each time step of the trajectory, a pseudorandom number generator was employed to determine step interval in x (left, right) and y (up, down) directions. Under these conditions, the diffusion coefficient, *D* is given by:

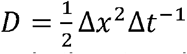

where ∆x is the step size and ∆t is the time interval. The grid size of the simulation was chosen to match the pixel resolution of the experimental microscope images (0.08678 μm/pixel). The time step was typically fixed at 0.16 seconds, however variation of this parameter within a factor of five had no effect on the results of the simulation. At each time step (n), the position of each point at step (n+1) was calculated and verified to lie within the nuclear boundary. If the new (n+1) position was determined to escape the nuclear region, reflecting boundary conditions were enforced by reverting the (n+1) position to the position at step (n). Trapping in ROI0 was similarly enforced. At each step (n), the point position was checked to determine whether it lay within the trapping region. If so, position at step (n+1) was set to the position at step (n). The trajectories of each of the 12,000 points were stored in memory for post-run analysis. Accumulation and depletion kinetics were determined by summing the total number of points in each ROI at each time step from the stored trajectories. The experimental data were fit empirically using r-squared coefficients between the simulated curves and the experimental data. We processed the accumulation kinetics for all the nuclei shown in Supp. Figure 4 – 7, and only analyzed selected depletion kinetics from regions adjacent to the DNA damage site. (Note that the depletion kinetics are accurately modeled only when the region-of-interest areas are located entirely inside the nucleus.)

